# A modular chromosomally integrated toolkit for ectopic gene expression in *Vibrio cholerae*

**DOI:** 10.1101/2020.03.02.973776

**Authors:** Triana N. Dalia, Jennifer L. Chlebek, Ankur B. Dalia

## Abstract

The ability to express genes ectopically in bacteria is essential for diverse academic and industrial applications. Two major considerations when utilizing regulated promoter systems for ectopic gene expression are (1) the ability to titrate gene expression by addition of an exogenous inducer and (2) the leakiness of the promoter element in the absence of the inducer. Here, we describe a modular chromosomally integrated platform for ectopic gene expression in *Vibrio cholerae*. We compare the broadly used promoter elements P_*tac*_ and P_*BAD*_ to versions that have an additional theophylline-responsive riboswitch (P_*tac*_-riboswitch and P_*BAD*_-riboswitch). These constructs all exhibited unimodal titratable induction of gene expression, however, max induction varied with P_*tac*_ > P_*BAD*_ > P_*BAD*_-riboswitch > P_*tac*_-riboswitch. We also developed a sensitive reporter system to quantify promoter leakiness and show that leakiness for P_*tac*_ > P_*tac*_-riboswitch > P_*BAD*_; while the newly developed P_*BAD*_-riboswitch exhibited no detectable leakiness. We demonstrate the utility of the tightly inducible P_*BAD*_-riboswitch construct using the dynamic activity of type IV competence pili in *V. cholerae* as a model system. The modular chromosomally integrated toolkit for ectopic gene expression described here should be valuable for the genetic study of *Vibrio cholerae* and could be adapted for use in other species.

## Introduction

Regulated promoter systems for ectopic gene expression have been widely used in bacterial systems. Two commonly employed system for ectopic gene expression are the isopropyl β-d-1-thiogalactopyranoside (IPTG)-inducible *tac* promoter (P_*tac*_) and the arabinose inducible *araBAD* promoter (P_*BAD*_) (1, 2). Both of these systems, however, exhibit some degree of leakiness, which allows for gene expression in the absence of the inducer when cells are grown in rich LB medium. Leakiness of P_*BAD*_ can be reduced, to some extent, by addition of glucose to the growth medium because this promoter is catabolite repressed (3), however, addition of glucose to the growth medium can change the physiology of cells which may introduce a confounding variable for some experiments.

Riboswitches are control elements that can regulate gene expression via direct interactions between a small molecule ligand and mRNA. Synthetic riboswitches that are responsive to the small molecule theophylline have recently been developed, which allow for regulated gene expression in diverse biological systems (4, 5). These riboswitch elements likely fold the mRNA to occlude the ribosome binding site in the absence of theophylline. And binding of theophylline to the riboswitch alters the conformation of the mRNA to expose the ribosome binding site and allow for translation of the downstream gene. For this reason, it is important to note that these riboswitches likely have limited utility for controlling the expression of genetic elements like non-coding RNAs, which do not need to be translated to exert their effect.

Generally, plasmids are employed for ectopic gene expression. However, many commonly used plasmids are poorly maintained by *Vibrio* species and/or their copy number can vary relative to model systems like *Escherichia coli* (6, 7). Integration of ectopic expression constructs onto the genome can bypass these issues. For many *Vibrio* species (e.g. *Vibrio cholerae, Vibrio natriegens, Vibrio campbellii, Vibrio vulnificus, Vibrio parahaemolyticus*, and *Vibrio fischeri*), it is remarkably easy to integrate novel sequences into the bacterial genome by exploiting their inherent capacity to undergo horizontal gene transfer by natural transformation (8-13), which can be exploited for ectopic gene expression (10, 13-17).

Here, we generate a chromosomally integrated modular platform for ectopic gene expression in *Vibrio* species based on the widely-used P_*tac*_ and P_*BAD*_ promoters in conjunction with a previously described theophylline responsive riboswitch (5). We demonstrate that this toolkit allows for differing levels of ectopic gene expression (i.e. max induction), and that one of these promoter constructs (P_*BAD*_-riboswitch) allows for a broad-range of titratable gene expression without detectable leakiness. We highlight the utility of this tight expression construct to study the dynamic surface appendages required for natural transformation in *V. cholerae*.

## Results & Discussion

### Design of modular ectopic expression constructs

All of the ectopic expression constructs are distinct ‘cassettes’ that can be integrated at any location in the bacterial genome (**Fig. 1**). We accomplish this via simple splicing by overlap extension (SOE) PCR (18) to stitch these expression cassettes to upstream and downstream regions of homology (see **Fig. S1** for details) to generate linear PCR products that can then be integrated into the *V. cholerae* genome by chitin-induced natural transformation (8). Once an expression construct is integrated at a genomic locus, the gene of interest to be ectopically expressed can be easily exchanged by SOE PCR and natural transformation (see **Fig. S1** for details). In this study, all ectopic expression constructs are integrated at the VCA0692 locus. This is a frame-shifted gene in the N16961 reference genome (19). Furthermore, the disruption of VCA0692 does not alter the fitness of *V. cholerae* during growth in rich medium or in environments this facultative pathogen encounters during its pathogenic life cycle (20), thus, highlighting this locus as a useful “neutral” location for the integration of novel sequences. These constructs can also be integrated at other commonly used “neutral” loci in *V. cholerae* including the *lacZ* gene, the frame-shifted transposase VC1807, or within intergenic spaces between convergently transcribed genes (21).

**Fig. 1.**
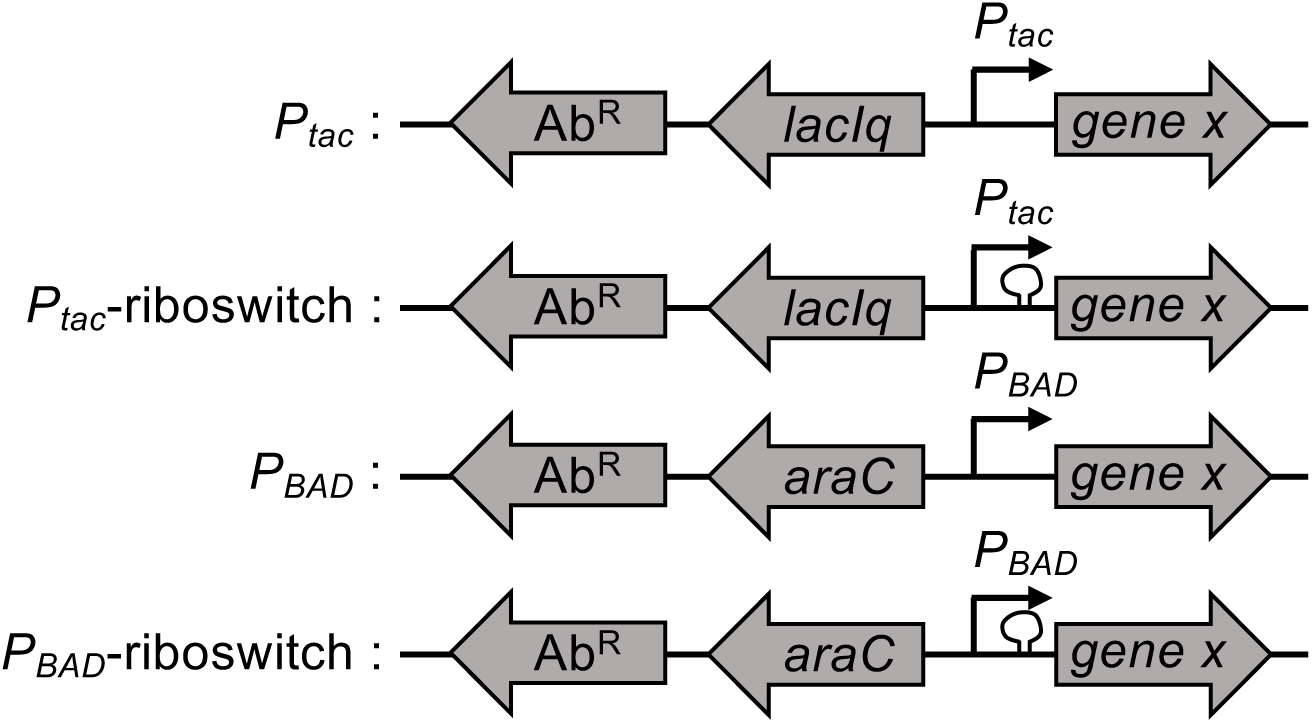
Diagram of ectopic expression constructs. The four ectopic expression constructs characterized in this study are indicated. All have a linked antibiotic resistance cassette (Ab^R^) to facilitate selection during integration into the genome by natural transformation. The gene encoding a transcription factor (*lacIq* or *araC*) and the promoter (P_*tac*_ or P_*BAD*_) required for inducible gene regulation are indicated. The presence of a theophylline-dependent riboswitch is demarcated by a loop before the gene of interest (*gene x*). For details on how these constructs were assembled, see **Fig. S1**.

The constructs for ectopic gene expression have a modular design where all have a linked antibiotic resistance marker (Ab^R^) (**Fig. 1**). This Ab^R^ facilitates selection during integration into the genome by natural transformation and can be easily altered depending on the need. Linked to this Ab^R^, there are the gene control elements. For P_*tac*_ and P_*tac*_-riboswitch constructs, this includes the LacIq repressor and *tac* promoter. By contrast the P_*BAD*_ and P_*BAD*_-riboswitch constructs contain AraC and the *araBAD* promoter. Both the P_*tac*_ and P_*BAD*_ promoter constructs can be engineered to have a user-defined ribosome binding site (**Fig. S1**). By contrast, the two riboswitch constructs (P_*tac*_-riboswitch and P_*BAD*_-riboswitch) contain a defined ribosome binding site within the theophylline-dependent riboswitch (riboswitch “E” in (5)) that is located immediately upstream of the gene of interest (**Fig. 1** and **Fig. S1**).

### Testing inducibility of ectopic expression constructs with GFP

To test whether these different chromosomally integrated constructs allow for inducible gene expression and to compare the maximum level of expression they support, we generated constructs for ectopic expression of *gfp* (22). The maximum level of gene expression varied among constructs with P_*tac*_ > P_*BAD*_ > P_*BAD*_-riboswitch > P_*tac*_-riboswitch (**Fig. 2**). Also, all of these constructs allowed for titratable gene expression (**Fig. 2**). This is particularly notable for P_*BAD*_, because this inducible system is known to have an “all-or-none” or autocatalytic gene expression phenotype in wildtype strains of *E. coli* (23). This autocatalytic expression profile is due to high affinity transport of arabinose in *E. coli*, which further stimulates increased expression of arabinose transporters. Uncoupling this autoregulatory loop in *E. coli* can allow for titratable gene expression (24). *V. cholerae* does not catabolize arabinose and lacks high affinity arabinose transporters. Arabinose may be transported into *V. cholerae* nonspecifically through one (or more) of its other carbohydrate transporters. Regardless, this low affinity transport of arabinose in *V. cholerae* allows for titratable gene expression from P_*BAD*_ (**Fig. 2**), which is consistent with a number of previous studies (14, 25). Also, for constructs that contained the additional riboswitch control element (P_*tac*_-riboswitch and P_*BAD*_-riboswitch), ectopic gene expression was dependent on addition of theophylline (**Fig. 2**), as expected (5).

**Fig. 2.**
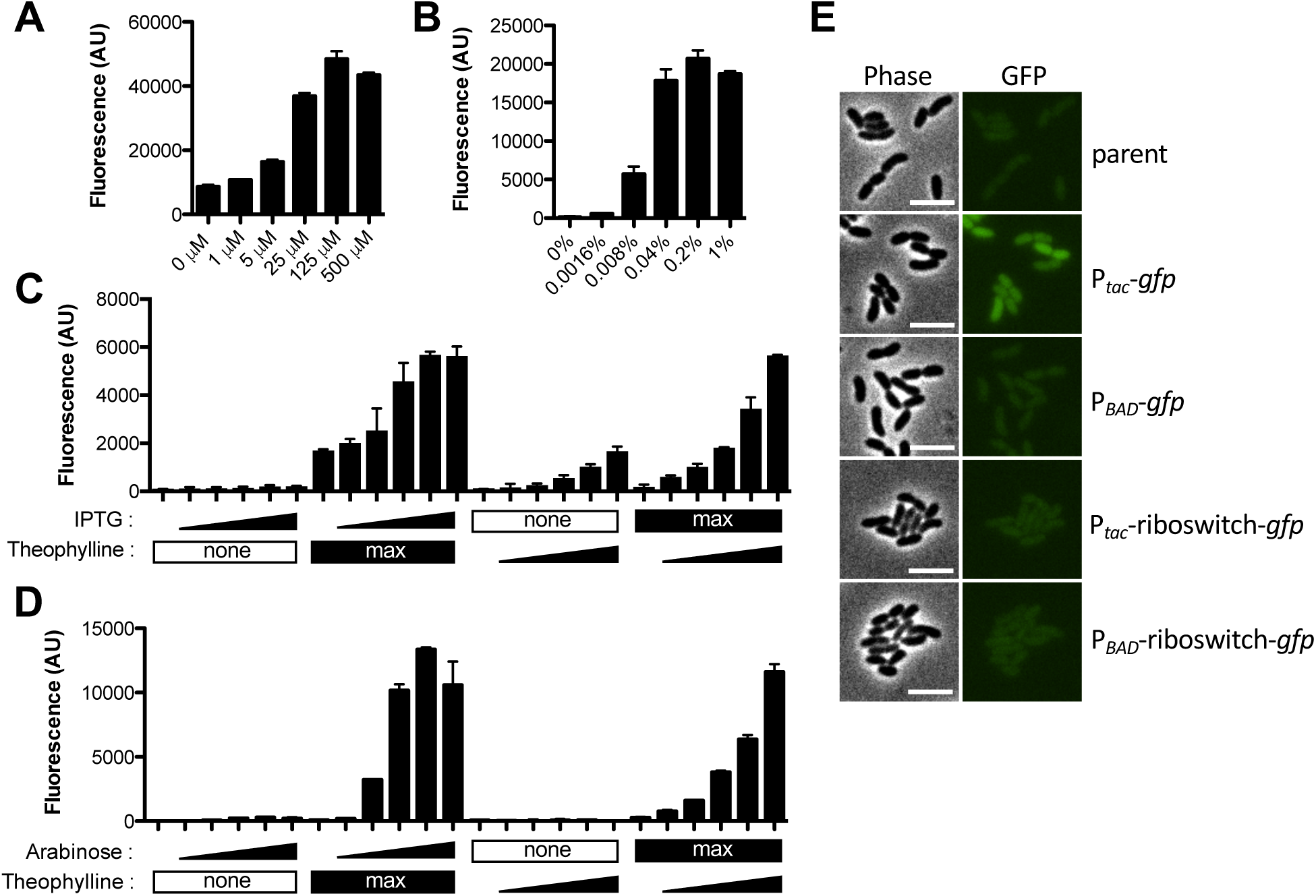
Testing inducibility of ectopic expression constructs with GFP. (**A**-**D**) Cells harboring ectopic expression constructs driving *gfp* integrated at the VCA0692 locus were grown with inducer as indicated and assessed for GFP fluorescence in bulk cultures on a plate reader. (**A**) Cells with a P_*tac*_-*gfp* construct were grown with the indicated amount of IPTG. (**B**) Cells with a P_*BAD*_-*gfp* construct were grown with the indicated amount of arabinose. (**C**) Cells with a P_*tac*_-riboswitch construct were grown with increasing doses (denoted by a triangle below bars) of IPTG (from left to right: 0 µM, 1 µM, 5 µM, 25 µM, 125 µM, 500 µM) or theophylline (from left to right: 0 mM, 0.018 mM, 0.054 mM, 0.16 mM, 0.5 mM, 1.5mM). ‘Max’ below bars denotes that cells were incubated with the highest concentration of IPTG (500 µM) or theophylline (1.5 mM) as indicated, while ‘none’ indicates that none of that inducer was added. (**D**) Cells with a P_*BAD*_-riboswitch construct were grown with increasing doses (denoted by a triangle below bars) of arabinose (from left to right: 0%, 0.0016%, 0.008%, 0.04%, 0.2%, 1%) or theophylline (from left to right: 0 mM, 0.018 mM, 0.054 mM, 0.16 mM, 0.5 mM, 1.5mM). ‘Max’ below bars denotes that cells were incubated with the highest concentration of arabinose (1%) or theophylline (1.5 mM) as indicated, while ‘none’ indicates that none of that inducer was added. All data in **A**-**D** are from at least two independent biological replicates and shown as the mean ± SD. (**E**) Representative phase and epifluorescence images of cells with the indicated ectopic expression construct grown without any inducer added. Scale bar, 4 µM.

### Testing ectopic expression constructs for the distribution of GFP fluorescence within single cells

While we observed titratable gene expression above (**Fig. 2**), this was assessed in bulk cultures. Thus, titratable gene expression could be the result of bimodality in gene expression where cells in the population exhibit either a highly fluorescent or poorly fluorescent phenotype (similar to an ON/OFF light switch); and increased inducer simply results in a shift within the population where a higher proportion of cells exhibit the highly fluorescent phenotype. This is in contrast to titratable gene expression where the population responds uniformly to yield a unimodal distribution where increased inducer simply shifts the fluorescence intensity of the entire population (similar to a dimmer switch). Generally, for ectopic expression constructs the latter phenotype is preferred. To distinguish between these possibilities, we assessed the distribution of fluorescence among single cells within induced populations by epifluorescence microscopy. In the absence of inducer, only the P_*tac*_ construct exhibited detectable GFP fluorescence (**Fig. 2A and E**), which is consistent with this construct being very leaky in *V. cholerae*; a phenotype that is already widely appreciated. In the presence of inducer, all four constructs exhibited unimodal distributions, which supports the latter model and suggests that there is a uniform response to inducer among single cells within the population (**Fig. 3**).

**Fig. 3.**
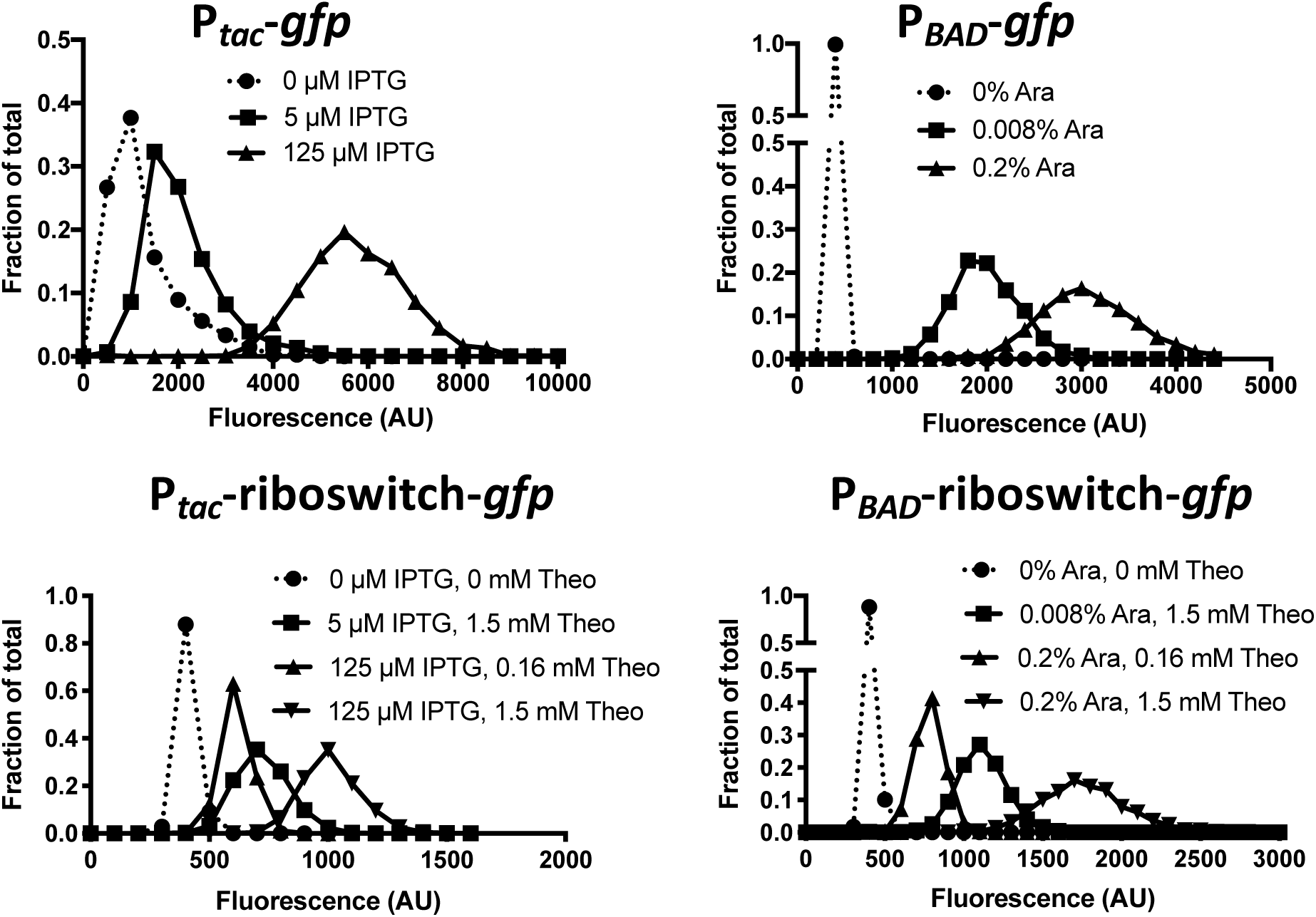
Testing ectopic expression constructs for the distribution of GFP fluorescence within single cells. Cells harboring the indicated ectopic expression constructs integrated at the VCA0692 locus were grown with inducer as indicated and assessed for GFP fluorescence in single cells via epifluorescence microscopy (Theo = theophylline; Ara = arabinose). The distribution of fluorescence among cells in the population is indicated on the plotted histograms. Data are from >1000 cells per condition tested and representative of two independent experiments.

### Testing leakiness of ectopic expression constructs with Flp recombinase

A major consideration for ectopic expression constructs is leakiness, which is defined as the basal expression of regulated genes in the absence of inducer. As mentioned above, only the P_*tac*_ construct exhibited detectable leakiness when using GFP fluorescence as a readout. This, however, is a poor readout for leaky gene expression because a substantial amount of GFP protein is required to generate an observable fluorescent readout. We sought to develop a sensitive and easily measured phenotype for leakiness from our ectopic expression constructs. To that end, we employed flippase (Flp), a highly-efficient recombinase that can mediate site-specific recombination between two Flp recombinase target (FRT) sequences (26, 27). Flanking FRT sequences can be engineered to leave behind an in-frame scar following Flp excision (28). To generate a simple readout for Flp-mediated activity, we introduced a FRT-flanked Ab^R^ into the *lacZ* gene in *V. cholerae* (**Fig. 4A**). Strains with *lacZ*::FRT-Ab^R^-FRT yielded a white colony phenotype on X-gal plates. Following Flp-mediated excision, however, the resulting *lacZ* gene (containing an in-frame 81 bp insertion) is active, and strains harboring *lacZ*::FRT exhibit a blue colony phenotype on X-gal plates (**Fig. 4A**). Thus, following Flp-mediated resolution, cells are irreversibly converted from LacZ-(white colonies) to LacZ+ (blue colonies). To determine whether our expression constructs exhibited leaky expression, we generated ectopic expression constructs to drive Flp expression in strains that harbored the *lacZ*::FRT-Ab^R^-FRT construct. P_*tac*_-*Flp* was so leaky that all cells where we introduced this construct resolved *lacZ*::FRT-Ab^R^-FRT to yield only blue colonies even in the absence of inducer. All of the other constructs (P_*tac*_-riboswitch-*Flp*, P_*BAD*_-*Flp*, and P_*BAD*_-riboswitch-*Flp*) yielded only resolved blue colonies when grown in the presence of inducer. In the absence of inducer, both P_*tac*_-riboswitch-*Flp* and P_*BAD*_-*Flp* exhibited some degree of leakiness, while P_*BAD*_-riboswitch-*Flp* did not exhibit any detectable leakiness in this assay (**Fig. 4B**). It is notable that the limit of detection of this assay is ∼3 logs below the leakiness observed from the P_*BAD*_ promoter, a construct that is traditionally considered tightly repressed in the absence of inducer (2, 3). This further validates our Flp-based assay as a highly sensitive approach to assess promoter leakiness.

**Fig. 4.**
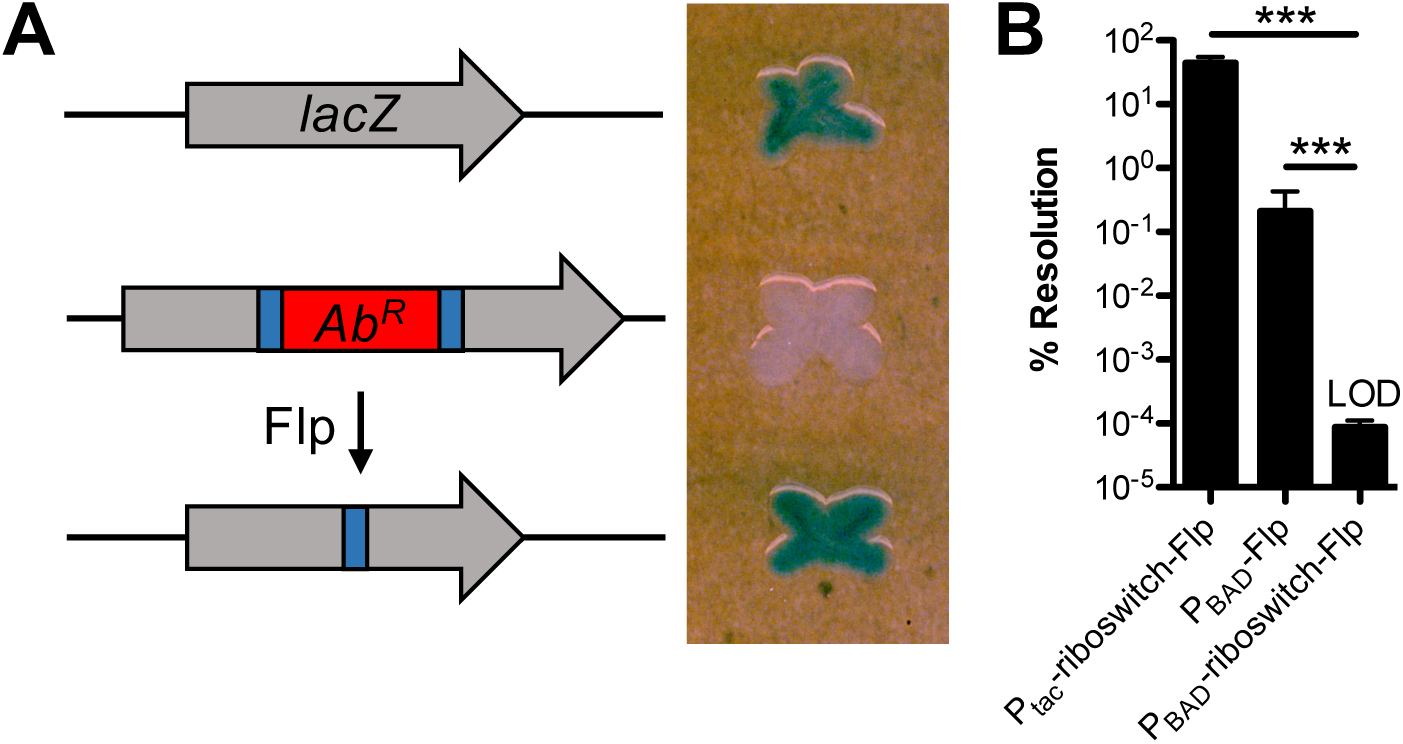
Testing leakiness of ectopic expression constructs with Flp recombinase. (**A**) Diagram of the approach used to test leakiness in gene expression with Flp recombinase. Wildtype *V. cholerae* cells have intact *lacZ* and form blue colonies on X-gal plates (top). Cells with *lacZ*::FRT-Ab^R^-FRT have inactive LacZ and are white on X-gal plates (middle). Flp recombination resolves the FRT-Ab^R^-FRT cassette within *lacZ* (making *lacZ*::FRT), which restores LacZ activity (bottom). (**B**) Single colonies of cells harboring the indicated ectopic expression constructs integrated at the VCA0692 locus and *lacZ*::FRT-Ab^R^-FRT were grown in LB medium without any inducer overnight. Then, % resolution of *lacZ*::FRT-Ab^R^-FRT was determined by plating for quantitative culture on X-gal plates. Percent resolution is defined as the number of blue colonies / total CFUs. Data in **B** are from 6 independent biological replicates and shown as the mean ± SD. Statistical comparisons were made by one-way ANOVA with Tukey’s post-test. LOD, limit of detection, *** = *P* < 0.001.

### Employing ectopic expression constructs to study the dynamic activity of type IV competence pili in V. cholerae

Horizontal gene transfer by natural transformation in *V. cholerae* is dependent on type IV competence pili (29, 30). These pili extend from the bacterial surface, bind to DNA in the environment, and then retract to pull DNA across the outer membrane (29). This ingested DNA can then be translocated into the cytoplasm and integrated into the bacterial genome by homologous recombination. PilB is the motor ATPase that is required for extension of type IV competence pili (31). To study the role of PilB in dynamic pilus activity, we sought to establish a strain that allowed for titratable and tightly regulated control of pilus extension. To that end, we generated strains where the native copy of *pilB* was deleted, and *pilB* expression was ectopically driven by our expression constructs. We then tested whether these strains were naturally transformable when grown without any inducer added. Only leaky expression of *pilB* would allow for natural transformation in this assay because Δ*pilB* mutants are not transformable (**Fig. 5A**). As observed using our Flp reporter readout, in the absence of inducer, P_*tac*_, P_*tac*_-riboswitch, and P_*BAD*_ all exhibited leaky expression of *pilB* as evidenced by detectable natural transformation, while there was no detectable leakiness observed for P_*BAD*_-riboswitch because no transformants were obtained in this background without inducer added (**Fig. 5A**). Importantly, this experiment was performed in plain LB medium, thus, induction of catabolite repression (by the addition of glucose to the medium) was not necessary to prevent leaky gene expression. As expected, all strains transformed at high rates in the presence of inducer (**Fig. 5A**). These results suggest that leaky expression of *pilB* from P_*tac*_, P_*BAD*_, and P_*tac*_-riboswitch allow for some degree of pilus assembly even without inducer added, while pilus assembly is completely inhibited in the P_*BAD*_-riboswitch construct when no inducer is present. To test this idea further, we deleted the retraction ATPase *pilT* in these strains. This prevents extended pili from being easily retracted and sensitizes the direct observation of pilus assembly via a recently developed pilus labeling approach (32). To determine whether leaky *pilB* expression allowed for pilus assembly, we assessed piliation in strains that were grown in the absence of inducer. Indeed, for P_*tac*_, P_*BAD*_, and P_*tac*_-riboswitch, we observed cells that contained extended pilus fibers (**Fig. 5B**), which is consistent with the leaky expression detected using our Flp recombinase reporter (**Fig. 4B**). We did not, however, observe extended pili in strains with P_*BAD*_-riboswitch when grown without inducer (**Fig. 5B**), which is consistent with a lack of leaky expression for this construct (**Fig. 4B**). As expected, all strains generated extended pili when grown with the appropriate inducer (**Fig. 5B**).

**Fig. 5.**
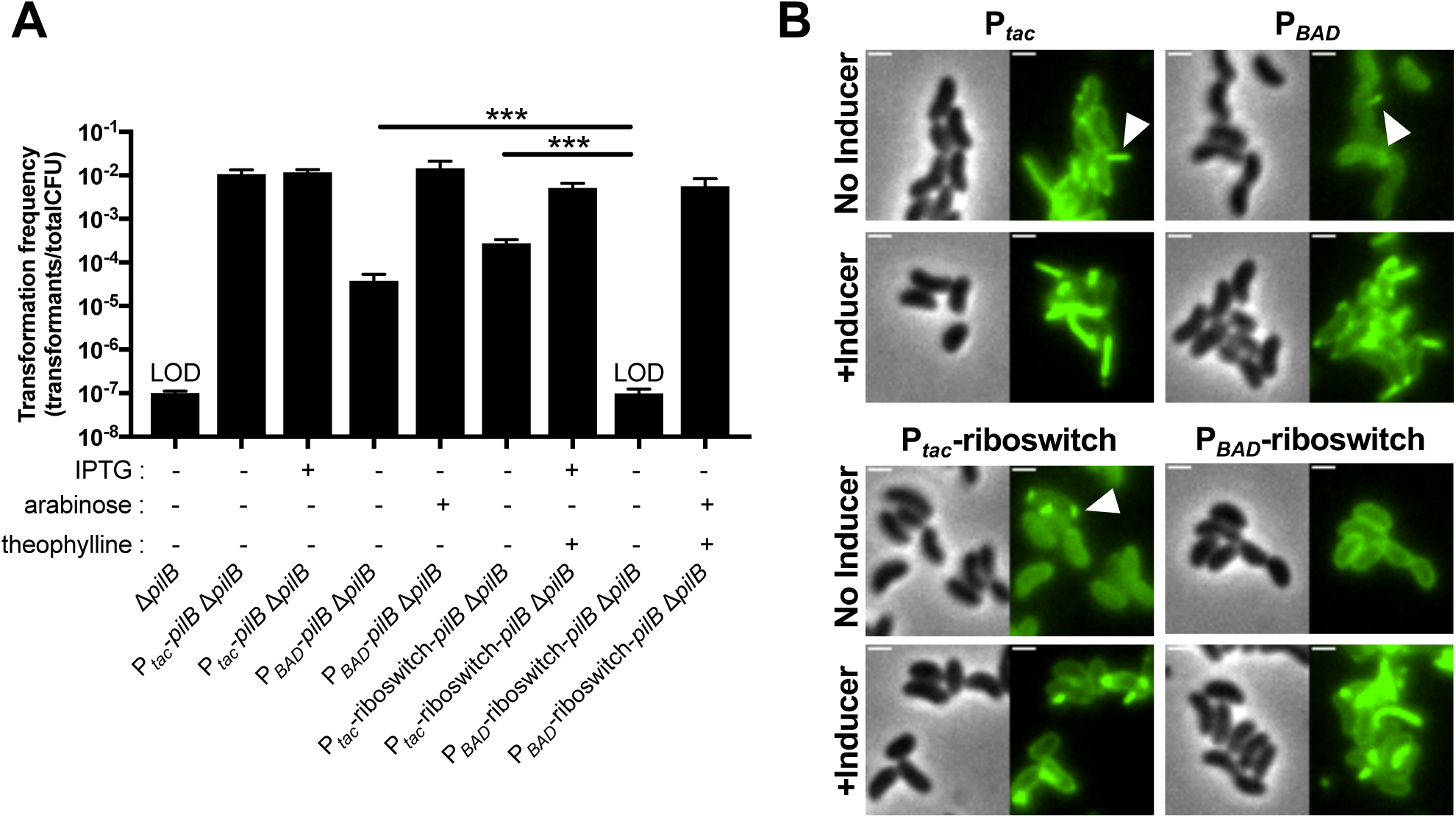
Employing ectopic expression constructs to study the type IV competence pilus extension ATPase PilB. (**A**) Natural transformation of the indicated strains was tested. All ectopic expression constructs in these strains were integrated at the VCA0692 locus. IPTG (100 µM), arabinose (0.2%), and/or theophylline (1.5 mM) were added to reactions as indicated below each bar. Data are from at least 3 independent biological replicates and shown as the mean ± SD. Statistical comparisons were made by one-way ANOVA with Tukey’s post-test. LOD, limit of detection, *** = *P* < 0.001. (**B**) Representative images of surface piliation. All strains harbor Δ*pilB* and Δ*pilT* mutations at the native locus and the indicated ectopic *pilB* expression construct. Where indicated, cells were grown with inducer as follows: 100 µM IPTG for P_*tac*_, 0.2% arabinose for P_*BAD*_, 100 µM IPTG + 1.5 mM theophylline for P_*tac*_-riboswitch, and 0.2% arabinose + 1.5 mM theophylline for P_*BAD*_-riboswitch. Examples of extended pili in no inducer samples are indicated by white arrows. Scale bars, 1 µm.

Together, these data indicate that only our newly generated P_*BAD*_-riboswitch-*pilB* construct allows for tightly regulated and titratable control of pilus biogenesis / extension in *V. cholerae*. This provides a valuable resource that will be critical for addressing diverse questions related to type IV pilus biology, which will be the focus of future work.

## Methods

### Bacterial strains and growth conditions

All strains used in this study are derivates of E7946, an El Tor isolate of *V. cholerae* (33). See **Table S1** for a complete list of strains used in this study. Strains were routinely grown at 30°C or 37°C in LB Miller broth and agar (BD Difco). When appropriate, media was supplemented with carbenicillin (20 µg/mL), spectinomycin (200 µg/mL), kanamycin (50 µg/mL), trimethoprim (10 µg/mL), chloramphenicol (1 µg/mL), or erythromycin (10 µg/mL).

### Construction of mutant strains

All strains were generated by SOE PCR and chitin-induced natural transformation exactly as previously described (8, 21). See **Fig. S1** for details on how the ectopic expression constructs were assembled and **Table S2** for a detailed list of all primers used to generate all of the mutant constructs in this study. The *araC* – P_*BAD*_ region for our P_*BAD*_ and P_*BAD*_-riboswitch constructs was amplified from pBAD18-Kan (3). The P_*tac*_-riboswitch construct was amplified from DNA generously provided by Kim Seed (15).

### GFP fluorescence assays

Cells harboring ectopic expression constructs driving *gfp* were grown rolling at 30°C to mid-log in LB medium. Then, inducer was added as indicated in each experiment and cells were grown for two additional hours rolling at 30°C. Next, cells were washed and resuspended to the same optical density in instant ocean medium (7 g/L; Aquarium Systems) and fluorescence was measured on a Biotek H1M plate reader: excitation 500 nm / emission 540 nm exactly as previously described (34). The parent strain lacking any ectopic expression construct was assayed alongside and used to subtract the background fluorescence of cells.

To image cells for GFP fluorescence, cultures were grown rolling at 30°C in LB medium in the presence of the indicated inducers for 5 hours to late-log. Then, cells were washed in instant ocean medium and mounted on 0.2% gelzan pads made with instant ocean medium exactly as previously described (35).

### Flp recombinase assays

Cells harboring ectopic expression constructs driving *Flp* and *lacZ*::FRT-Spec^R^-FRT were struck out onto LB + X-gal (40 µg/mL) + spectinomycin (200 µg/mL) plates. Single white colonies were picked, inoculated into plain LB medium, and grown overnight rolling at 30°C. Then, each culture was plated quantitatively on LB+Xgal plates to determine the frequency of blue colonies within the population. Empirically, we determined that we could only reliably detect blue colonies at a rate of ∼1 in 1,000,000 cells or 0.0001%. This equated to scoring for blue colonies on 100 µL spread plates of a dilution of 10^−3^ or greater. If no blue colonies were observed at 10^−3^, we assumed a single blue colony was present to define the limit of detection for that sample. Data are reported as the % Resolution = (CFU/mL blue colonies / CFU/mL total colonies) x 100.

### Natural transformation assays

Cells harboring ectopic expression constructs driving *pilB* and the native *pilB* gene deleted were tested for rates of natural transformation. All of these strains also harbored P_*constitutive*_-*tfoX* and Δ*luxO* mutations, which rendered these strains constitutively competent. Strains were tested for natural transformation using chitin-independent transformation assays exactly as previously described (36). The transforming DNA using in these experiments was 100 ng of a 6 kb ΔVC1807::Erm^R^ PCR product.

### Pilus labeling

All strains harbored the indicated ectopic expression construct, a cysteine amino acid substitution in the major pilin subunit PilA (PilA^S56C^), a deletion of the native copy of *pilB* (Δ*pilB*), P_*constitutive*_-*tfoX*, a Δ*luxO* mutation, and a deletion of the retraction ATPase *pilT* (Δ*pilT*). Strains were grown and labeled with Alexa fluor 488-maleimide exactly as previously described (36). And mounted on 0.2% gelzan pads made with instant ocean medium.

### Microscopy

Phase contrast and fluorescence images were collected on a Nikon Ti-2 microscope using a Plan Apo ×60 objective, a GFP filter cube, a Hamamatsu ORCAFlash 4.0 camera and Nikon NIS Elements imaging software. For **Fig. 2E**, the lookup tables for each phase and fluorescent image were adjusted to the same range so that they can be compared between samples. Fluorescence intensity of cells was determined using the MicrobeJ plugin (37) in Fiji (38).

## Acknowledgements

We gratefully acknowledge Eric Bruger, Christopher Marx, Julia van Kessel, and Johann Strnat for helpful discussions; and Kim Seed for generously providing DNA with the P_*tac*_-riboswitch construct. This work was supported by funds from the National Science Foundation (1714949), Department of Energy (DE-SC0019436), and the National Institutes of Health (R35GM128674) to ABD.

## Supporting Information

**Fig. S1.**
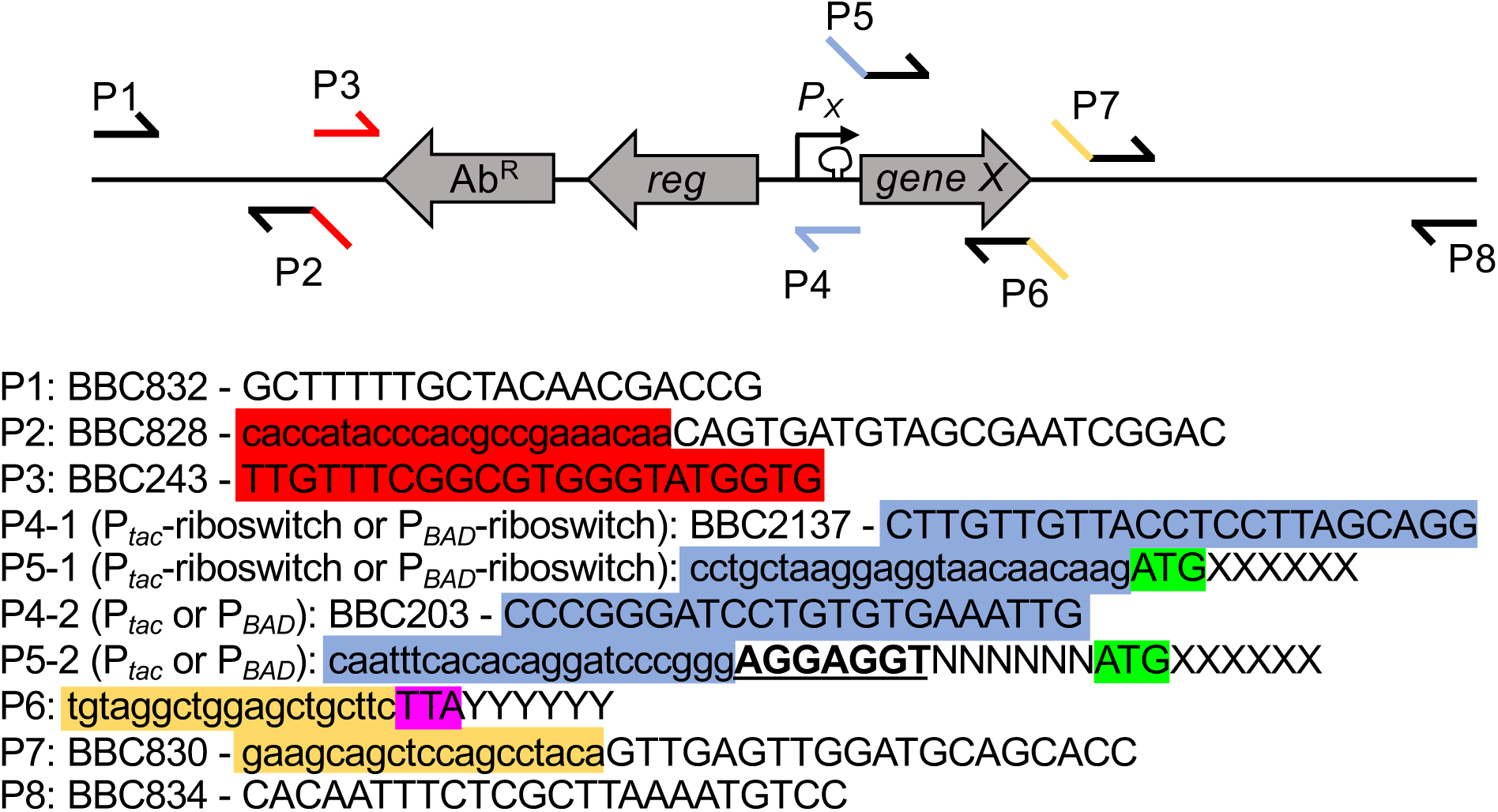
Details on how to assemble the ectopic expression constructs and swap out genes of interest. A generic diagram of the ectopic expression construct is shown with the location of different primers needed to initially construct each expression construct (P1, P2, P3, P6, P7, and P8). As well as the primers to insert novel genes of interest into established expression constructs (P1, P4, P5, P6, P7, and P8). Ab^R^ = antibiotic resistance cassette, reg = regulatory gene (*lacIq* or *araC*), P_X_ = the promoter (P_*tac*_, P_*BAD*_, P_*tac*_-riboswitch, or P_*BAD*_-riboswitch), and gene X = the gene of interest.

### Establishing ectopic expression constructs at new genomic locations

The UP arm of homology is generated with P1 and P2, while the DOWN arm of homology is amplified with P7 and P8. In this example, these primers amplify homology arms to integrate the ectopic expression constructs into *V. cholerae* ChII. P2 and P7 contain tails that overlap any of the 4 ectopic expression constructs. To move intact expression constructs to new locations in the genome, they can be amplified with P3 and P6. This can serve as a MIDDLE arm to stitch to the UP and DOWN arms of homology.

### To insert new genes of interest

Once an ectopic expression construct has been introduced to a genetic locus, the gene of interest can be swapped out.

-To amplify a gene of interest to place within P_*tac*_-riboswitch or P_*BAD*_-riboswitch the forward P5-1 primer must have a 5’ overlap region as indicated. Immediately after this overlap should be the start codon for the gene of interest (highlighted in green) + additional sequence to serve as a primer for the gene of interest (denoted by ‘XXXXXX’). P5-1 overlaps with P4-1 (blue highlighted sequence), which sits within the theophylline-dependent riboswitch. Thus, the same overlap can be used to make either P_*tac*_-riboswitch or P_*BAD*_-riboswitch constructs. The reverse P6 primer for the gene of interest must have the tail indicated, which should be immediately followed by the stop codon of the gene (highlighted in purple – TAA stop codon in this example) + additional sequence to serve as a primer for the gene of interest (denoted by ‘YYYYYY’).

-To amplify a gene of interest to place within P_*tac*_ or P_*BAD*_ the forward P5-2 primer must have the 5’ overlap region indicated. Immediately after this overlap should be the desired ribosome binding site (indicated in bold and underline – the ‘optimal’ RBS AGGAGGT is used in this example), which should be followed by a 6 bp spacer (this spacing between the ribosome binding site and the start codon is essential for optimal translation and can be derived from the native gene) + the start codon (highlighted in green) + additional sequence to serve as a primer for the gene of interest (denoted by ‘XXXXXX’). P5-2 overlaps with P4-2 (blue highlighted sequence), which sits downstream of the two promoter elements. Thus, the same overlap can be used to make either P_*tac*_ or P_*BAD*_ constructs. The reverse P6 primer for the gene of interest can be made exactly as described above.

All amplified genes of interest serve as MIDDLE arms in SOE reactions with an UP arm amplified with P1 and P4, and a DOWN arm amplified with P7 and P8.

**Table S1.** Strains used in this study

| Name | Relevant figure(s) | Full genotype | Reference / Strain# |
| --- | --- | --- | --- |
| parent | Fig. 2E | E7946 *lacZ*::FRT-Spec<sup>R</sup>-FRT | This study / SAD035 |
| *P<sub>tac</sub>-gfp* | Fig. 2A, E; Fig. 3 | E7946 *lacZ*::FRT-Spec<sup>R</sup>-FRT, ΔVCA0692::*P<sub>tac</sub>-gfp* Carb<sup>R</sup> | This study / TND2290 (SAD2780) |
| *P<sub>BAD</sub>-gfp* | Fig. 2B, E; Fig. 3 | E7946 *lacZ*::FRT-Spec<sup>R</sup>-FRT, ΔVCA0692::*P<sub>BAD</sub>-gfp* Carb<sup>R</sup> | This study / TND2291 (SAD2781) |
| *P<sub>tac</sub>-riboswitch-gfp* | Fig. 2C, E; Fig. 3 | E7946 *lacZ*::FRT-Spec<sup>R</sup>-FRT, ΔVCA0692::*P<sub>tac</sub>-riboswitch-gfp* Carb<sup>R</sup> | This study / TND2293 (SAD2782) |
| *P<sub>BAD</sub>-riboswitch-gfp* | Fig. 2D, E; Fig. 3 | E7946 *lacZ*::FRT-Spec<sup>R</sup>-FRT, ΔVCA0692::*P<sub>BAD</sub>-riboswitch-gfp* Carb<sup>R</sup> | This study / TND2292 (SAD2783) |
| wildtype | Fig. 4A | E7946 | (25) / SAD031 |
| *P<sub>BAD</sub>-Flp* | Fig. 4A-B | E7946 *lacZ*::FRT-Spec<sup>R</sup>-FRT, ΔVCA0692::*P<sub>BAD</sub>-Flp* Carb<sup>R</sup> | This study / TND2083 (SAD2784) |
| *P<sub>tac</sub>-riboswitch-Flp* | Fig. 4B | E7946 *lacZ*::FRT-Spec<sup>R</sup>-FRT, ΔVCA0692::*P<sub>tac</sub>-riboswitch-Flp* Carb<sup>R</sup> | This study / TND2289 (SAD2785) |
| *P<sub>BAD</sub>-riboswitch-Flp* | Fig. 4B | E7946 *lacZ*::FRT-Spec<sup>R</sup>-FRT, ΔVCA0692::*P<sub>BAD</sub>-riboswitch-Flp* Carb<sup>R</sup> | This study / TND2086 (SAD2786) |
| Δ*pilB* | Fig. 5A | E7946 Sm<sup>R</sup>, *P<sub>const</sub>*-tfoX, ΔluxO, *pilA* S67C, *lacZ*::FRT-Kan<sup>R</sup>-FRT, ΔVC1807::*CmR*, Δ*pilB* | This study / TND2373 (SAD2787) |
| *P<sub>tac</sub>-pilB* Δ*pilB* | Fig. 5A | E7946 Sm<sup>R</sup>, *P<sub>const</sub>*-tfoX, ΔluxO, *pilA* S67C, Δ*lacZ*::*P<sub>tac</sub>-pilB* Spec<sup>R</sup>, ΔVC1807::*CmR*, Δ*pilB* | This study / JLC769 (SAD2788) |
| *P<sub>BAD</sub>-pilB* Δ*pilB* | Fig. 5A | E7946 Sm<sup>R</sup>, *P<sub>const</sub>*-tfoX, ΔluxO, *pilA* S67C, *lacZ*::FRT-Kan<sup>R</sup>-FRT, ΔVC1807::*CmR*, Δ*pilB*, ΔVCA0692::*P<sub>BAD</sub>-pilB* Carb<sup>R</sup> | This study / TND2403 (SAD2789) |
| *P<sub>tac</sub>-riboswitch-pilB* Δ*pilB* | Fig. 5A | E7946 Sm<sup>R</sup>, *P<sub>const</sub>*-tfoX, ΔluxO, *pilA* S67C, *lacZ*::FRT-Kan<sup>R</sup>-FRT, ΔVC1807::*CmR*, Δ*pilB*, ΔVCA0692::*P<sub>tac</sub>-riboswitch-pilB* Carb<sup>R</sup> | This study / TND2402 (SAD2790) |
| *P<sub>BAD</sub>-riboswitch-pilB* Δ*pilB* | Fig. 5A | E7946 Sm<sup>R</sup>, *P<sub>const</sub>*-tfoX, ΔluxO, *pilA* S67C, *lacZ*::FRT-Kan<sup>R</sup>-FRT, ΔVC1807::*CmR*, Δ*pilB*, ΔVCA0692::*P<sub>BAD</sub>-riboswitch-pilB* Carb<sup>R</sup> | This study / TND2378 (SAD2791) |
| *P<sub>tac</sub>-pilB* Δ*pilB* Δ*pilT* | Fig. 5B | E7946 Sm<sup>R</sup>, *P<sub>const</sub>*-tfoX, ΔluxO, *pilA* S67C, Δ*lacZ*::*P<sub>tac</sub>-pilB* Spec<sup>R</sup>, ΔVC1807::*CmR*, Δ*pilB*, Δ*pilT*::Zeo<sup>R</sup> | This study / TND2498 |
| *P<sub>BAD</sub>-pilB* Δ*pilB* Δ*pilT* | Fig. 5B | E7946 Sm<sup>R</sup>, *P<sub>const</sub>*-tfoX, ΔluxO, *pilA* S67C, *lacZ*::FRT-Kan<sup>R</sup>-FRT, ΔVC1807::*CmR*, Δ*pilB*, ΔVCA0692::*P<sub>BAD</sub>-pilB* Carb<sup>R</sup>, Δ*pilT*::Zeo<sup>R</sup> | This study / TND2501 |
| $P_{tac}$-riboswitch-*pilB*<br>$\Delta pilB \Delta pilT$ | Fig. 5B | E7946 Sm<sup>R</sup>, $P_{const}$-tfoX, $\Delta luxO$, *pilA* S67C,<br>*lacZ*::FRT-Kan<sup>R</sup>-FRT, $\Delta VC1807$::CmR, $\Delta pilB$,<br>$\Delta VCA0692$::$P_{tac}$-riboswitch-*pilB* Carb<sup>R</sup>,<br>$\Delta pilT$::Zeo<sup>R</sup> | This study /<br>TND2500 |
| $P_{BAD}$-riboswitch-*pilB*<br>$\Delta pilB \Delta pilT$ | Fig. 5B | E7946 Sm<sup>R</sup>, $P_{const}$-tfoX, $\Delta luxO$, *pilA* S67C,<br>*lacZ*::FRT-Kan<sup>R</sup>-FRT, $\Delta VC1807$::CmR, $\Delta pilB$,<br>$\Delta VCA0692$::$P_{BAD}$-riboswitch-*pilB* Carb<sup>R</sup>,<br>$\Delta pilT$::Zeo<sup>R</sup> | This study /<br>TND2499 |

**Table S2.**
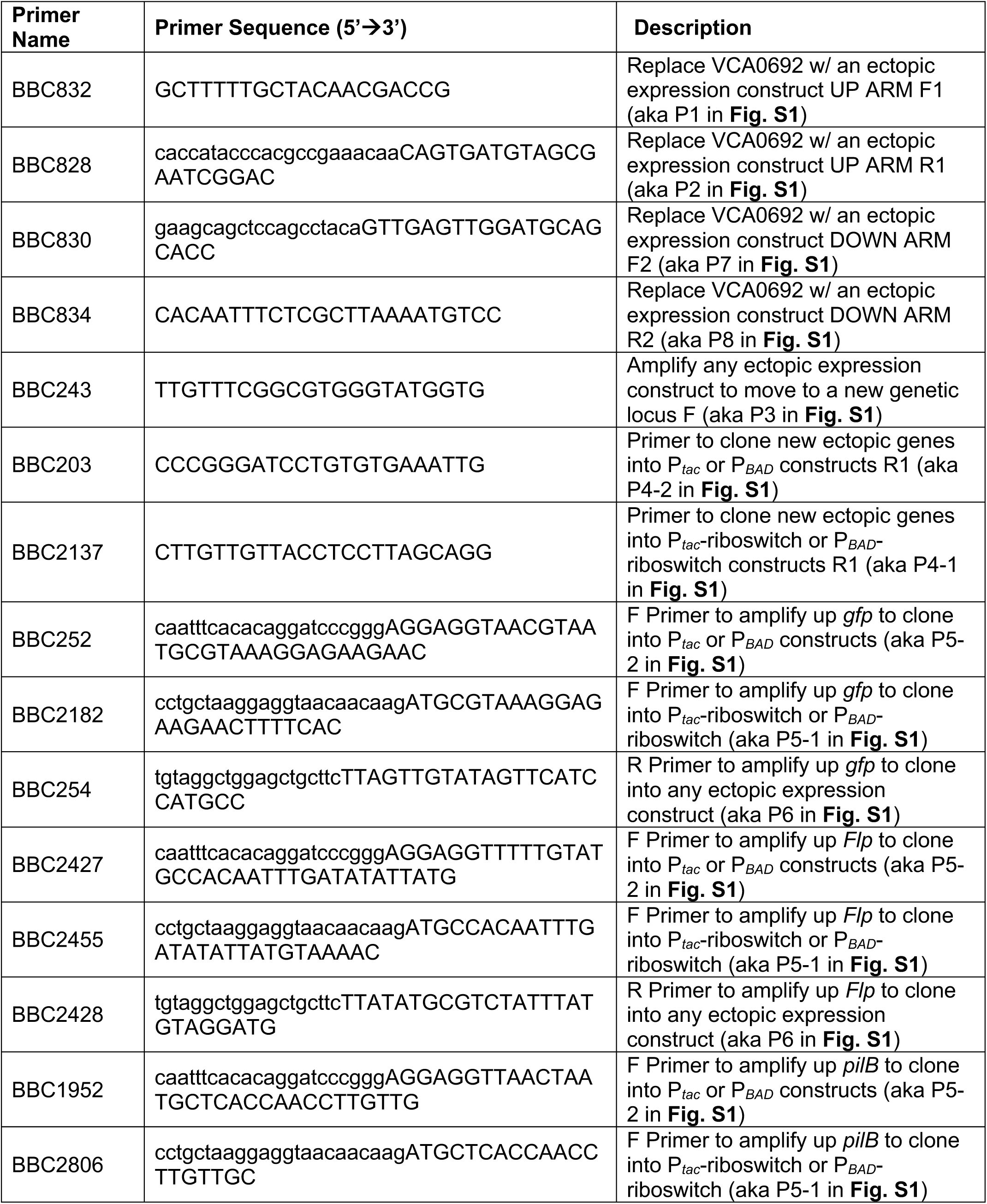
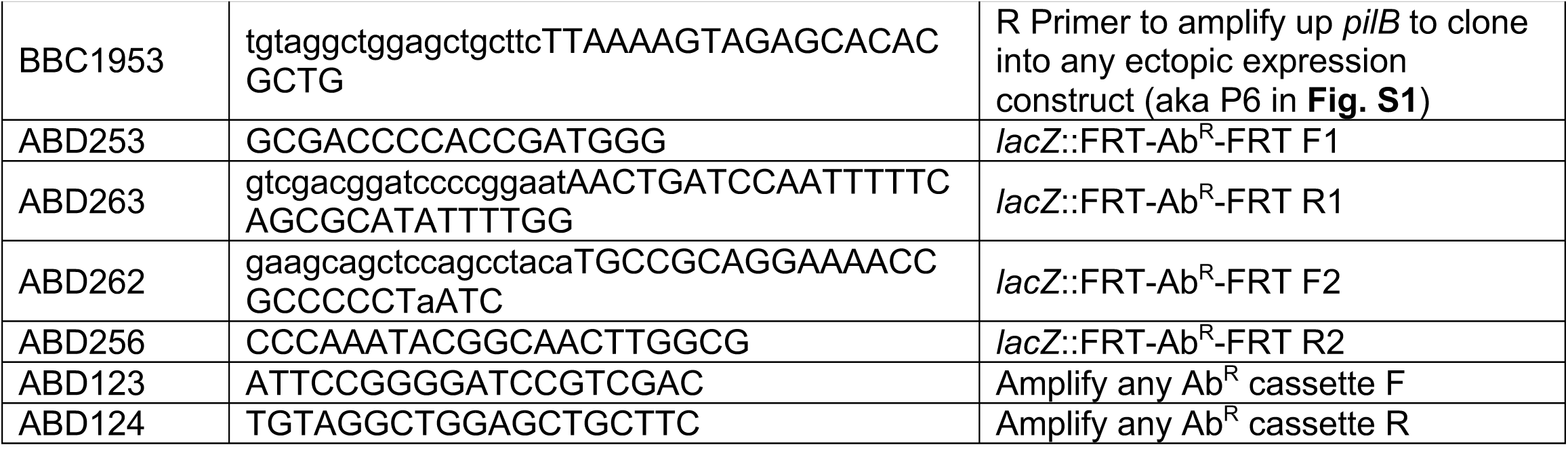
Primers used in this study

## Notes

### Competing Interest Statement

The authors have declared no competing interest.

